# Biodiversity effects on ecosystem functioning: disentangling the roles of biomass and effect trait expression

**DOI:** 10.64898/2026.03.17.711861

**Authors:** Alice N. Ardichvili, Markus Bittlingmaier, Grégoire T. Freschet, Michel Loreau, Jean-François Arnoldi

## Abstract

Species diversity potentially has a dual effect on communities: a generally positive effect on overall community biomass, reflecting the expression of species response and interaction traits, and a poorly characterised effect on mass-specific species contribution to ecosystem functions, reflecting the expression of their effect traits. Disentangling the effects of biodiversity on total biomass from those on effect trait expression would help settle a long-standing debate by clarifying how biodiversity relates to both facets of species effects on ecosystem functioning.
Following the classical BEF approach, we calculate expected ecosystem function based on observed functioning in monoculture. We then derive a net biodiversity effect (NBE) and decompose it into four components: the classical complementarity and selection effects on total community biomass, and complementarity and selection effects on effect trait expression. The latter two reflect, respectively, a complementarity or facilitation in how effect traits influence the function, and how species with the highest potential for increasing the function become dominant in the community.
We illustrate this NBE decomposition with three ecosystem functions (nitrogen retention capacity, soil hydraulic conductivity improvement, and forage digestibility) measured in assembled communities under controlled experimental conditions of perennial grassland plants. Regarding nitrogen retention, we find a positive complementary effect via total biomass, but a negative biodiversity effect via effect trait expression. For hydraulic conductivity improvement, biodiversity effects are mostly mediated by total biomass. As for forage digestibility, we found a positive complementarity effect on trait expression, outweighed however by a negative selection effect. This analysis reveals how biodiversity may have contrasting effects on ecosystem functions via its impact on biomass and effect trait expression.

**Synthesis:** Separating between the effect of biodiversity on plant community biomass and on effect trait expression at the community level is one important step towards understanding the pathways by which diverse plant communities drive ecosystem functioning.

## Motivation

The rapid decline of biodiversity undermines the ecosystem processes that sustain life (IPBES, 2019). Whether diversity itself has a role beyond that of its constituent species has important consequences, and has been the subject of several decades of research. While the role of biodiversity on biomass production in plant communities is well-established (Cardinale et al., 2007, Loreau et al., 2022), its role in ecosystem processes that rely on particular functional traits (and their expression in a community context) has largely been overlooked.

Despite their conceptual simplicity, interpretations of Biodiversity and Ecosystem Functioning (BEF) experiments have revealed multiple, interacting mechanisms through which biodiversity impacts ecosystem functioning (i.e., any ecosystem variable involved in biotic-abiotic interactions; cf. glossary). Diversity can represent a reservoir of potentially highly productive species (Tilman et al., 1997), provide insurance against fluctuating environments due to species-specific responses of different magnitudes (Loreau and De Mazancourt, 2013), and enhance resource use efficiency because coexisting species use resources complementarily (McKane et al., 2002; Loreau and Hector, 2001, Wan et al., 2025). However, BEF relationships are not universally positive, as demonstrated for plant productivity (Dee et al. 2023), litter decomposition (Porre et al. 2020), or chitin degradation in pitcher plants (Cuellar-Guempler et al., 2024). Negative BEF relationships occur particularly for narrow functions, i.e. functions that are performed only by a subset of species and for which species contributions are highly uneven, or when competitive hierarchies are strong (D’Andrea et al., 2024). In such cases, increasing diversity may reduce functioning by intensifying competition on the most effective species or by diluting their dominance. As the focus in current ecological literature shifts from single to multiple ecosystem variables (Orr et al., 2024, Guo et al., 2025), considering contributions to both narrow and broad functions becomes increasingly important. This may require to clearly disentangle the relative importance of biodiversity effects on total biomass from these on community-level effect trait expression.

The classical BEF approach quantifies biodiversity effects (synonymous of non-additive or synergistic effects) as deviations of mixture performance from monoculture expectations (Loreau and Hector, 2001). These deviations result either from the dominance of species with a high potential for the function considered, that is, a selection effect, or from a complementarity effect. Net biodiversity effects reflect the outcome of biotic interactions (among one group of organisms, e.g. plant–plant interactions): community performance is more than the sum of all of its constituent species’ performances.

### Glossary

**Ecosystem variable *V*:** a scalar measure made at the ecosystem level, including properties (e.g. soil pH), pools (soil carbon stock), fluxes (soil N leaching) and processes (atmospheric N-fixation). Synonymous of a measure of an ‘ecosystem function’ understood in the broad sense of Hooper et al. (2005), but here we restrict the term ‘function’ to designate the ‘role’ or ‘contribution’ of a community in determining the value of an ecosystem variable (see Figure 1).

**Biomass-scalable function *Φ*:** an extensive measure of the contribution of a community to an ecosystem variable *V*, in the precise sense that there exist a transformation *x* ↦ *f* (*x*), independent of the community, such that *V* = *f* (*Φ*), where *Φ* scales with community biomass. Certain ecosystem variables directly qualify as biomass-scalable functions (e.g. net primary productivity). But for variables such as such as inorganic element pools (soil inorganic N), which can be non-zero in the absence of the species, identifying *Φ* requires to find and invert the transformation *x* ↦ *f* (*x*), which can be non-linear. Intensive properties, independent of total biomass, do not qualify as biomass-scalable function (e.g. forage digestibility) but can qualify as a mass-specific community contribution (see below) .

**Traits:** phenotypic features measured at the individual level (Violle et al., 2007). They can be classified (non exclusively) in three categories, depending on whether they pertain to the way organisms affect ecosystem variables, respond to the environment, or interact with each other.

**Effect traits:** determine the *direct* impact of organisms on a focal ecosystem variable (e.g. traits determining litter decomposability, Lavorel and Garnier, 2002).

**Response traits:** pertain to the way organisms respond to changes in environmental conditions (e.g. traits influencing the capacity of plants to resist a drought).

**Interaction traits:** pertain to the way organisms interact (e.g. allopathy). By definition, community dynamics reflect both response and interaction traits. By this logic, interaction and response traits determine the way organisms *indirectly* impact a focal ecosystem variable by determining community total biomass and species relative abundance.

**Mass-specific species contribution** *c*_*i*_ : Expression at the species-level of effect traits that determines the contribution of a species to an ecosystem function, per unit of biomass. Synonymous of the *Specific Effect Function* of Diaz et al. (2013)

**Mass-specific community contribution** 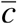 : Expression at the community-level of effect traits that determines the contribution of a community to an ecosystem function, per unit of biomass. Intensive property of the community, synonymous of the *Community Functional Parameter* of Violle et al. (2007), it is equal to the community-weighted mean of mass-specific species contributions.

**Net Biodiversity Effect (NBE)**: For a given function, the NBE is defined as the proportion of a biomass-scalable function that cannot be predicted from species monoculture biomass and mass-specific contributions to the function. In a very general sense, the NBE results from species interactions. Synonymous to *non-additivity* sensu Guo et al. (2025).

**Figure 1.**
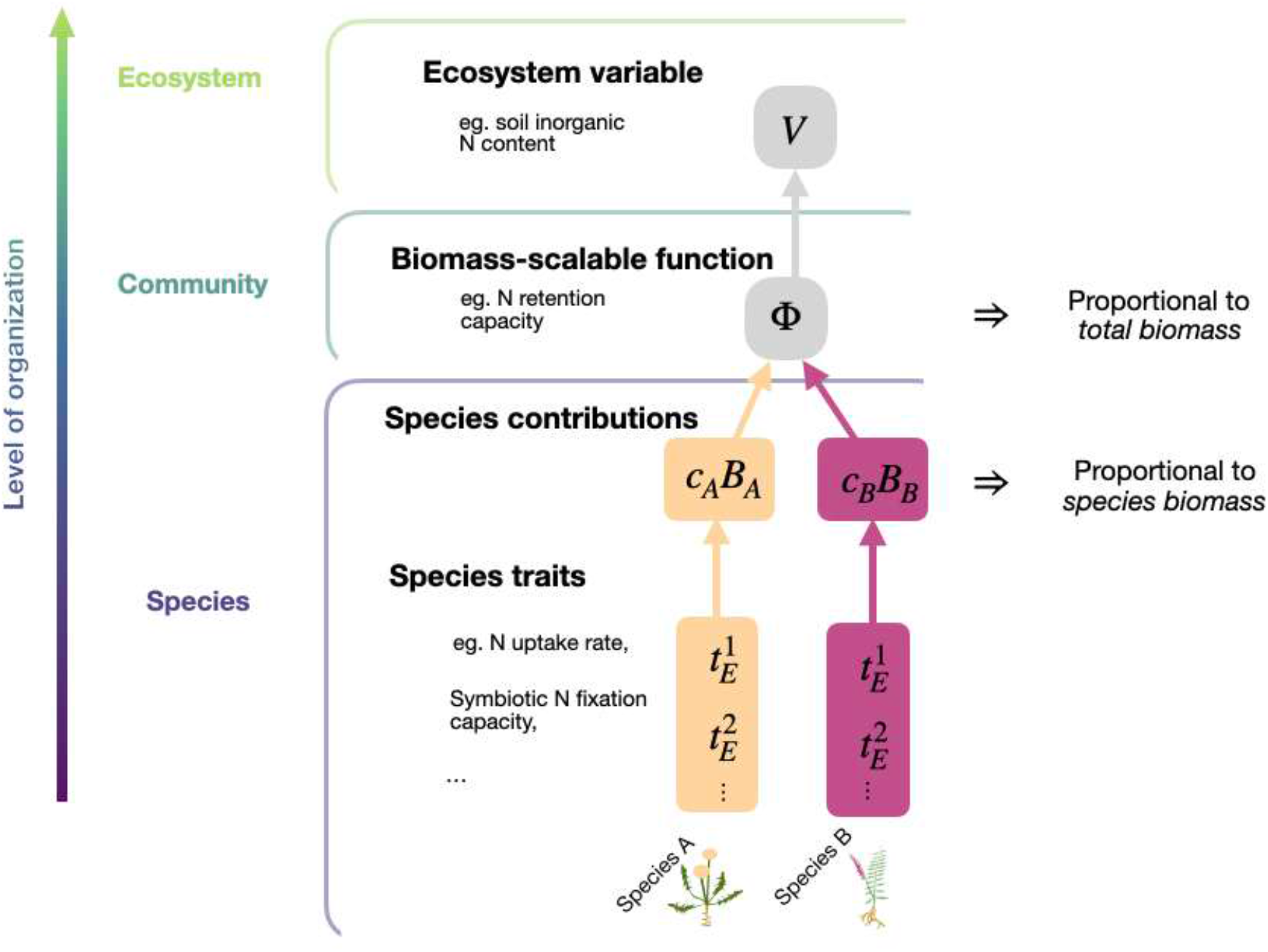
Scaling up from species-specific attributes to ecosystem variables. One or more species traits may determine their contribution, which is the product of species biomass and mass-specific contributions (see glossary). The biomass-scalable function describes the extent to which a community contributes to an ecosystem variable. It is the sum of its constituent species’ contributions and is therefore proportional to the community total biomass. How the biomass-scalable function relates to the ecosystem variable depends on our mechanistic understanding of the processes at play (see Box 1).

### Box 1: From ecosystem variables to biomass-scalable functions

In our framework, we analyse *biomass-scalable functions*, which denote a community’s contribution to the value of an ecosystem variable *V* (see Glossary). We then decompose the function according to the mass-specific contributions of species *c*_*i*_ and their biomass *B*_*i*_:

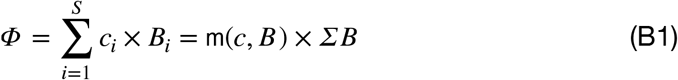

When there are no contributing organisms (no plants in our case), *ΣB* = 0 and the biomass-scalable function is null. By contrast, some ecosystem variables, such as soil N content or soil hydraulic conductivity, are not necessarily zero without plants. Without appropriate conversion, these variables therefore do not qualify as a biomass-scalable function under our definition. Such conversions can be achieved using simple mechanistic models, more comprehensive approaches like nutrient cycling models incorporating stoichiometry (Daufresne, 2021), in the vein of processes-informed metrics (PIMs, Levine et al., 2024), or phenomenological approaches. Below, we demonstrate this conversion from the ecosystem variable *soil inorganic N* to the biomass-scalable function *N retention capacity*.

We start with a simple nutrient cycling model (Loreau, 1998). Suppose that the inorganic Nitrogen content *N* in the soil solution of an experimental unit (in [mg]) follows standard chemostat dynamics:

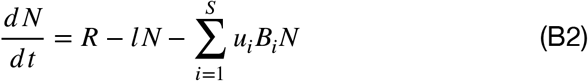

Where *R* [mg t^-1^] is the input rate, *lN* [mg t^-1^] is the leaching rate, *u*_*i*_ [mg^-1^ t^-1^] is the uptake rate of species *i* and *B*_*i*_ [mg] is the biomass of species *i*. At equilibrium, the N concentration in a pot without plants reads 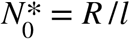 and the N concentration in a pot with plants reads 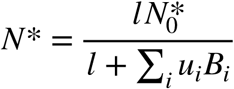 such that the community uptake rate reads: 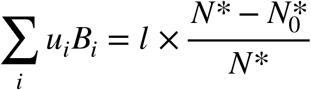

By dividing by the leaching rate, we obtain a measure of how uptake of inorganic N relates to its leaching rate, which we will term the N retention capacity:

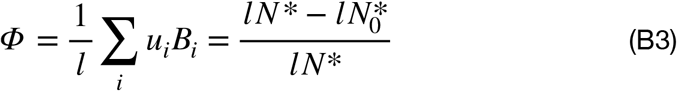

While the measurement of inorganic N content in the soil does not qualify as a biomass-scalable function, N retention capacity – which is proportional to the N uptake rate by a plant community – does. The N retention capacity can be readily obtained from (the difference in) measurements of leaching rates in the focal community and in bare soil.

While biodiversity effects on total biomass are well documented (Cardinale et al., 2007, Loreau et al., 2022), biodiversity effects on other ecosystem functions remains less understood. Biotic interactions can modify the expression of effect traits at the community level through changes in species relative abundance (Bruelheide et al., 2018). In addition, the value of a species’ effect traits may change in different communities due to plastic or micro-evolutionary responses (Biswas et al., 2025). The combination of several effect trait values may also result in synergistic effects. For example, the flammability of litter mixtures cannot be explained by the additive flammability of constituent species in isolation. It is the mixing of deciduous leaves with large surface areas and pines needles rich in flammable compounds that enhances air circulation and combustion through synergistic interactions independent of biomass (Magalhaes and Schwilk, 2012). Global changes may modify both biomass- and trait-mediated biodiversity effects by altering trait expression and species relative abundances (Vahsen et al., 2023, Wang et al., 2022). Disentangling additive from non-additive pathways in both total biomass and effect trait expression is therefore critical for understanding the role of biodiversity in driving ecosystem functions, predicting ecosystem responses to global change, and effectively conserving the eroding services biodiversity provides (Violle et al., 2007).

## Proposed framework

### From ecosystem variables to biomass-scalable functions

A key step in our approach consist in identifying a community’s « function », denoted *Φ*, that is, a community’s contribution to a given ecosystem-level variable of interest *V*, (see Glossary and Figure 1). Thus we assume that *V* = *f* (*Φ*) where the transformation *x* ↦ *f* (*x*) (which depends on the ecosystem variable, but not on the community of interest) is known, allowing us to quantify the function *Φ* from a measure of the variable *V* (see Box 1 for a worked out example). We further assume that *Φ*, all else being equal, is proportional to community biomass *ΣB*, seen as a proxy of the community’s potential to have an impact on the ecosystem (when *ΣB* = 0 there is no community, and thus it performs no function: *Φ* = 0). We thus assume that the function is an extensive property of species biomass. This first technical step is fundamental as it allows to build a sensible additive expectation for *Φ* based on monoculture observations.

In what follows, we focus on what we call « biomass-scalable functions », which can be written as 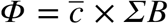 where 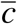 represents the « mass-specific contribution » of the community to the function (see Glossary). Crucially 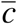 is assumed to not depend on total biomass, but could potentially depend on community composition, abiotic factors and so on. Note that 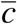 could be the notion of interest instead *Φ* of representing a *quality* of the community (e.g. forage digestibility instead of total humid matter of aboveground biomass, see below). In fact, our formalism aims to disentangle biodiversity effects on the *functional quality* vs. *quantity* of a given community.

### A mass-ratio representation of biomass-scalable functions

Any biomass-scalable function *Φ* can be seen as the sum of every species « mass-specific contributions » to the function *c*_*i*_ (see Glossary) multiplied by its biomass *B*_*i*_ (Eq. 1). Alternatively, the biomass-scalable function can be thought of as the product of total community biomass and the community-weighted mean of mass-specific species contributions (see glossary). In a community comprised of S species:

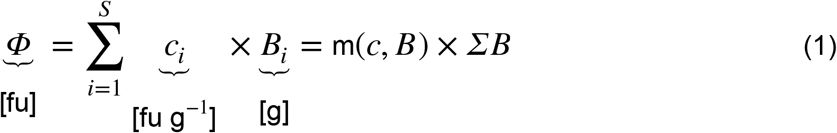

where m(*c, B*) denotes the average over mass-specific species contributions *c*_*i*_ of each species, weighted by their relative biomass, and gives the mass-specific community contribution to the function, so 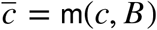 (see Glossary). In the rest of the article, we denote m(*x, w*) the average over the set of values *x* = {*x*_*i*_; *i* = 1, …, *S*} weighted by the set of non-negative values 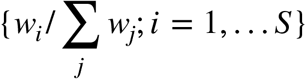. When all weights are equal (*w*_*i*_ ≡ 1), we omit the reference to the weights and write m(*x*) instead of *m*(*x*, 1), which is the classic arithmetic mean of *x*.

For example, if the biomass-scalable function is total plant biomass, then *c*_*i*_ = 1 for all species. If the biomass-scalable function is root biomass, the mass-specific species contribution *c*_*i*_ is the root mass fraction of each species. For some biomass-scalable functions, such as N retention capacity, the mass-specific effect *c*_*i*_ integrates several plant effect traits (Diaz 2025) relating, for instance, to the plant’s ability to take up N along the soil profile, retain it in its tissues and influence soil properties (Moreau et al., 2019; Freschet et al., 2021).

Equation 1 represents a ‘mass-ratio’ model of ecosystem functions consistent with Grime’s mass-ratio hypothesis, which posits that species contribute to ecosystem functioning in proportion to their relative biomass (Grime et al., 1998, Fig. 1). Box 1 provides guidance on whether any given ecosystem variable qualifies as a biomass-scalable function, and if it does not, how to define a biomass-scalable function from it.

### Obtaining the expected functioning based on monoculture observations

As in classical partitioning approaches (Loreau and Hector 2001), we quantify biodiversity effects by deriving expected polyculture functioning based on monoculture observations. In a monoculture of species *j*, we can measure the biomass-scalable function 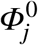 (in [fu]), and the biomass of species *j* in isolation *K*_*j*_ (in g). Applying Eq. 1 to a monoculture, we have:

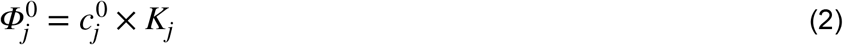

From the measures of 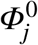 and *K*_*j*_, we can infer the mass-specific contribution of species *j* in the monoculture: 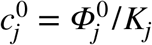 (in [fu] per g).

We now turn to polycultures. If species are planted in proportions 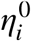, and assuming neutral effects among species, the expected biomass of species *i* is 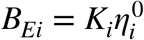 (as in Loreau and Hector 2001). Assuming further that mass-specific species contributions do not change with the biotic context, so that 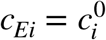, the expected functioning in the polyculture *Φ*_*E*_ becomes:

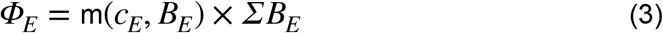

Where m(*c*_*E*_, *B*_*E*_) is the expected mass-specific community contribution. On the other hand, expected total biomass reads

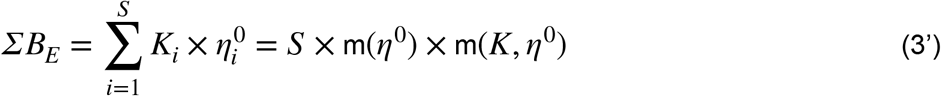

Since m(*η*^0^) = 1/*S*, and as in Loreau and Hector (2001), expected total biomass is just m(*K, η*^0^), the average biomass in monoculture (i.e. carrying capacity, *K*) weighted by the seeded densities.

### Decomposing the Net Biodiversity Effect

For a given community, we define the Net Biodiversity Effect (NBE) on the function *Φ* as the ratio of observed function *Φ*, to expected function, *Φ*_*E*_ (Eq. 3). When the observed function equates the expected one, the NBE is equal to one (Fig. 2A), and we detect no overall diversity effect. Note that previous decompositions (e.g. Loreau and Hector 2001) define the NBE as a difference between *Φ* and *Φ*_*E*_. Using a ratio makes the NBE a relative notion without units, thus easier to compare across systems. As it turns out, it also makes the following decomposition in various effects much simpler. This ratio reads:

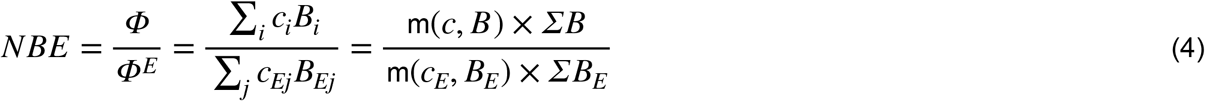

**Figure 2.**
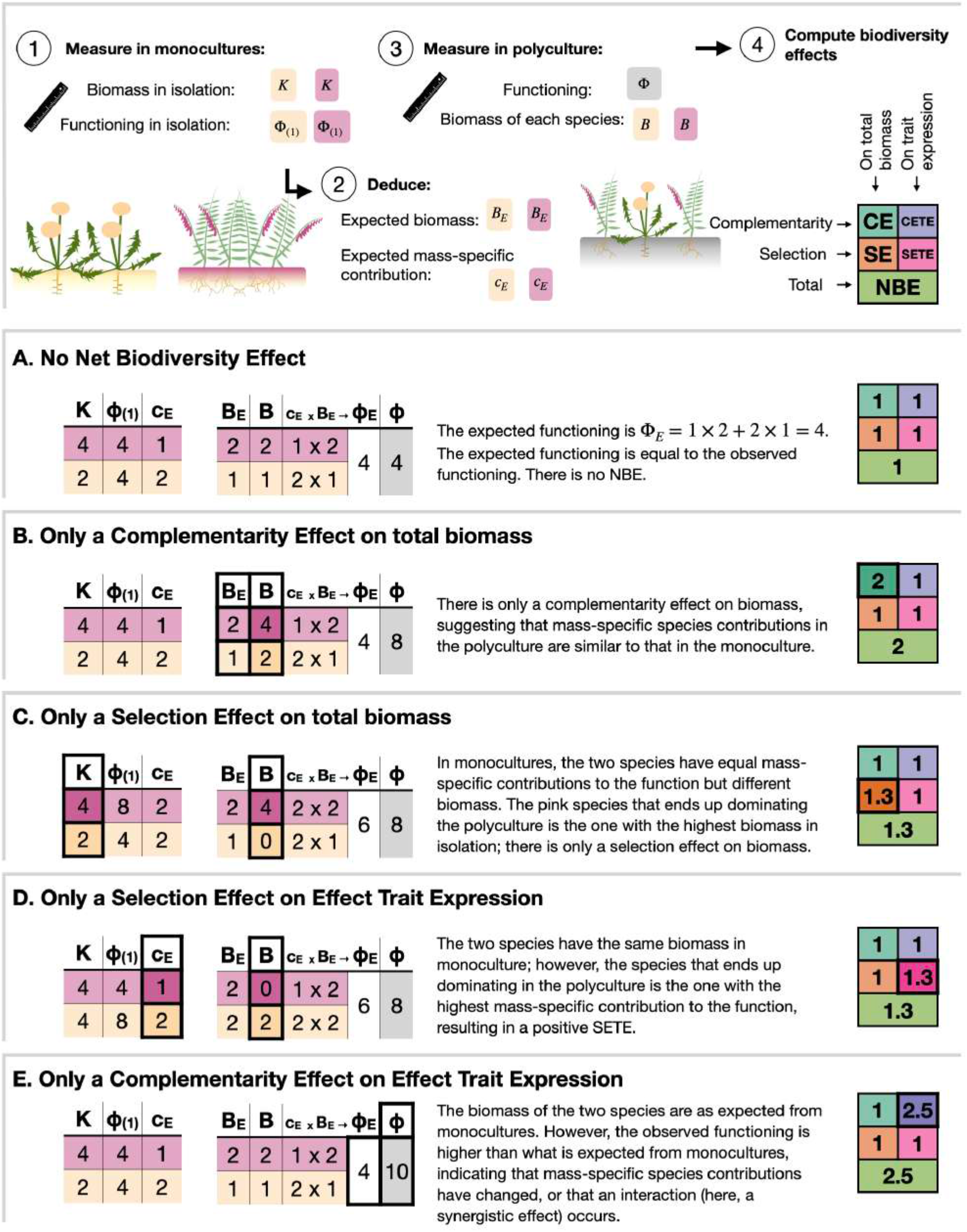
Different outcomes of a two-species polyculture, illustrating extreme cases in which only one component of the NBE, out of the four proposed in this partitioning, drives the positive NBE.

We denote *η*_*i*_ = *B*_*i*_ /*K*_*i*_ as the relative yield of species *i*, capturing the extent to which species *i* is favoured or suppressed in polyculture relative to monoculture (Vandermeer, 1989). Note that the expected relative yield *η*_*Ei*_ = *B*_*Ei*_ /*K*_*i*_ is simply 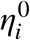, the seeded proportion of species i. We then rewrite total biomass as:

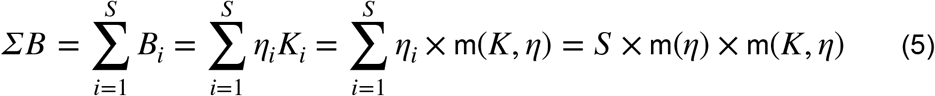

such that the ratio of total biomass to expected biomass, using Eq 3’, becomes:

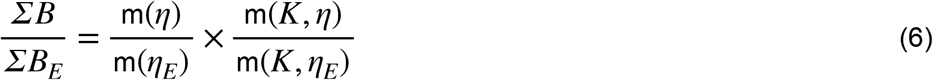

Plugging Eq. 6 into Eq. 4, and then introducing the average m(*c*_*E*_, *B*) to control for the effect of ecological assembly on the mass-specific community contribution, we obtain the following decomposition of the ratio of observed to expected functioning:

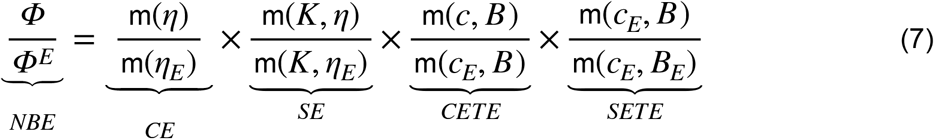

Equation 7 reveals the different components of the NBE, which fall in two categories. On the one hand, m(*K, η*)/m(*K, η*_*E*_) and m(*c*_*E*_, *B*)/m(*c*_*E*_, *B*_*E*_) are ratios of *averages of the same quantities* (*K* and *c*_*E*_, respectively), but *weighted differently* (using observed weights in the nominator and expected weights in the denominator). The difference between these weights captures a selection effect, i.e., how the community leads to dominance of species with high potential according to their monoculture performance (Fig.2 C-D). This potential can either reflect biomass production (Selection Effect on total biomass; SE) or mass-specific contributions arising from the expression of effect traits (Selection Effect on Effect Trait Expression; SETE). On the other hand, m(*η*)/m(*η*_*E*_) and m(*c, B*)/m(*c*_*E*_, *B*) are ratios of averages that are w*eighted by the same proportions*, but involve two different quantities. These contrast an expectation based on monocultures with a realisation, and thus capture a *complementarity effect*, in the sense that the community realisation is different from the sum of its non-interacting parts (Fig. 2B-E). As before, this difference can either reflect biomass production (Complementarity Effect on total biomass; CE) or mass-specific contributions (Complementarity Effect on Effect Trait Expression; CETE). Note that taking the log of Eq. 6 would give an additive partitioning of the NBE.

The complementarity effect on biomass, *CE* = m(*η*)/m(*η*_*E*_), simplifies to 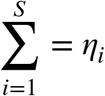, the sum of relative yields. Only when this sum (also termed *Land Equivalent Ratio* or *Relative Yield Total* in other studies; Vandermeer 1989) exceeds 1 can transgressive over-yielding occur, that is a greater community biomass than the highest monoculture biomass. From Eq. 1, m(*c, B*) = *Φ* /*ΣB*, such that the complementarity effect on trait expression, *CETE* = m(*c, B*)/m(*c*_*E*_, *B*), can be evaluated empirically even when the mass-specific contribution of species in polycultures cannot be directly assessed:

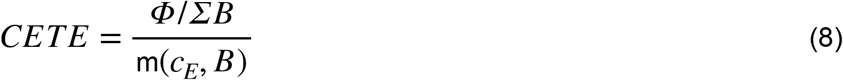

### Application to biomass-scalable functions performed by perennial plant communities

To illustrate our decomposition, we use data from an experiment quantifying the effect of perennial plant communities on eight ecosystem variables (Bittlingmaier *et al*., 2026), focusing here on three ecosystem variables: forage digestibility, N leachate concentration and soil hydraulic conductivity. Eight perennial grassland species were grown in either monocultures or in six-species polycultures (28 communities in total) in a greenhouse for 15 months.

In each plot, several variables were measured, including species-specific above-ground biomass in [g], plot-level total biomass *B*, N leachate concentration [*N*] in [mg L^-1^], humid matter content in [mg g^-1^] of plant aboveground biomass, and soil hydraulic conductivity *SHC* in [mm h^-1^]. The experimental set-up and methodology used to obtain these ecosystem variables are fully described in Bittlingmaier et al. (2026).

From these ecosystem variables, we computed two biomass-scalable functions (see glossary): N retention capacity and soil hydraulic conductivity *improvement*. Following the theoretical development in Box 1, N retention capacity was computed in each plot *u* as:

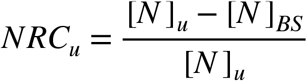

Where [*N* ]_*BS*_ and [*N* ]_*u*_ denote the concentration of N in leachates in bare-soil controls (averaged over pots without plants) and in planted pots, respectively. Soil hydraulic conductivity improvement quantifies how plants’ presence alters hydraulic conductivity (with positive values indicating improvement, which was the case for most observations) relative to the bare soil control. Soil hydraulic conductivity improvement was computed as:

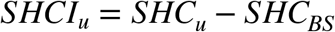

where *SHC*_*BS*_ and *SHC*_*u*_ denote soil hydraulic conductivity in bare soil controls and in planted pots, respectively.

Finally, forage digestibility being a *quality* rather than a quantity; it does not qualify as a biomass-scalable function. However, it can be approximated by the community-level trait humid matter content (Bumb 2018, Muraina 2025), which has units [mg g^-1^]. As such, humid matter content is a mass-specific contribution. The associated biomass-scalable function would be: the humid content of aboveground biomass, but the latter bears no agro-ecological meaning. Therefore, for *forage digestibility*, we computed only the biodiversity effects on effect trait expression, decomposed into complementarity and selection effects on effect trait expression (CETE and SETE, respectively). For all functions considered, we obtained the expected mass-specific species contributions 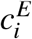 from the average of four monoculture replicates per species. We then computed the associated NBE and its different components for all six-species polycultures based on species biomass and observed functioning in each plot, according to Eq. 7.

## Application

Biodiversity effects differed strongly among the studied functions (Fig. 3), ranging from no net effect on N retention capacity (NBE = 1) to clearly positive effects on soil hydraulic conductivity (NBE > 1). Applying the presented partitioning framework demonstrates how biodiversity effects can be decomposed into its effect on ‘interaction trait’ expression driving community total biomass and its effect on ecosystem function via ‘effect trait’ expression, thereby revealing underlying and potentially counteracting mechanisms. This distinction between interaction traits and effect traits is intended as an analytical partition separating pathways affecting community biomass from those linking biomass to ecosystem function; some traits may therefore contribute to both pathways. Because complementarity and selection effects on total biomass were computed with the same expected vs. observed biomasses for soil hydraulic conductivity improvement and N retention capacity, the CE and SE are the same on both N retention capacity and soil hydraulic conductivity (compare Fig. 3a-b; turquoise and orange box plots are of the same magnitude —note the change of scales). However, for N retention capacity these biomass-driven gains were partly offset by the negative impact of effect trait expression: both CETE and SETE were negative. Overall, this indicates that, in species-rich communities, the average mass-specific species contribution to N retention is lower than expected from monocultures. More specifically, the negative SETE indicates that species with comparatively low mass-specific contribution to N retention tended to become dominant in polycultures. The negative CETE further suggests that, in species-rich communities, adjustments in effect trait values led to less positive (or more negative) impacts of species on the function. Regarding soil hydraulic conductivity, biodiversity effects mediated by effect trait expression were highly variable, showing neither consistently positive nor negative patterns. This variability points to a strong dependency of diversity effects on the community composition, potentially driven by species-specific neighbour interactions.

**Figure 3.**
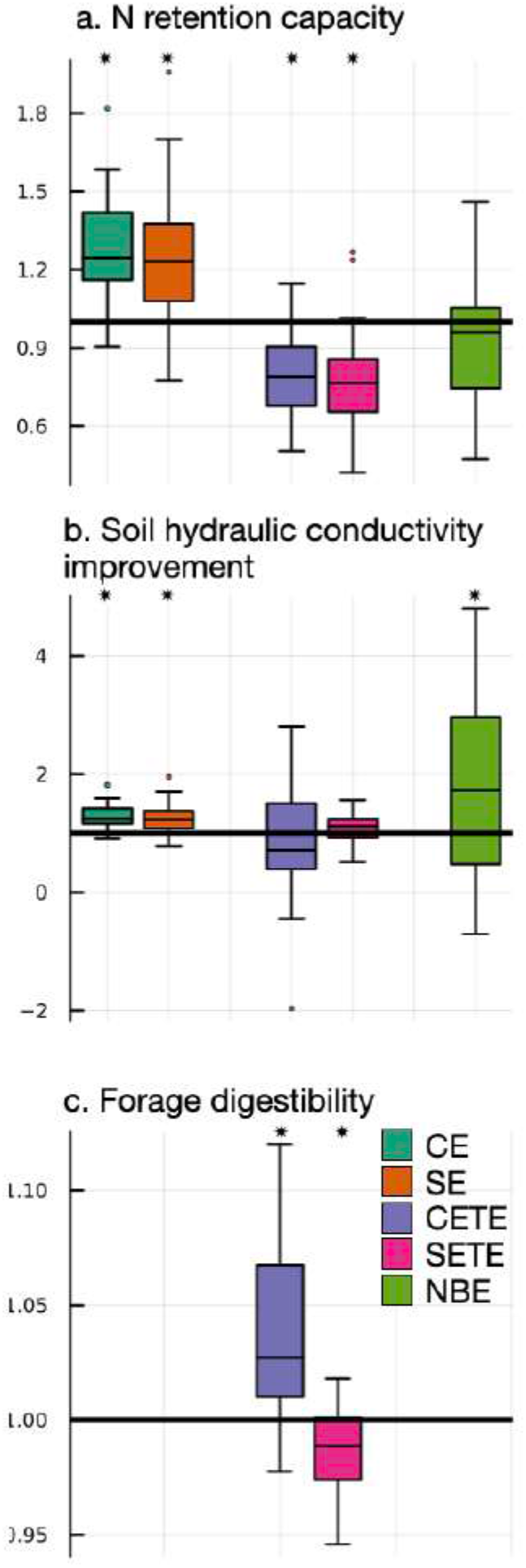
Decomposition of the Net Biodiversity Effect (NBE) into Complementarity Effect on total biomass (CE), Selection Effect on total biomass (SE), Complementarity Effect on Effect Trait Expression (CETE), and Selection Effect on Effect Trait Expression (SETE). The framework is illustrated using two biomass-scalable functions and one mass-specific community contribution, measured in six-species communities under controlled conditions: nitrogen (N) retention capacity (a), soil hydraulic conductivity improvement (b), and forage digestibility (c). Asterisks indicate that the average biodiversity effect differs significantly from 1 (p ≤ 0.05).

Forage digestibility provides a third, complementary illustration of the framework’s utility (Fig. 3c). Because this function is independent of total biomass, partitioning biodiversity effects into total biomass components (CE and SE) is not meaningful. Instead, it can be partitioned into trait-expression components. In doing so, we detected strong positive complementarity (CETE > 1), indicating that species richness enhances the digestibility of harvested biomass via adjustments of effect traits that affect digestibility. At the same time, a negative selection effect (SETE < 1) reveals that community dynamics favoured species with relatively low forage digestibility.

Together, these examples illustrate how the proposed partitioning disentangles the effect of biodiversity *via* total biomass – co-driven by the expression of ‘interaction traits’ – and the effect of biodiversity *via* the expression of effect traits. By separating these components, the method exposes opposing pathways that would otherwise remain hidden in net biodiversity effects, thereby allowing for a more nuanced, more mechanistic interpretation of biodiversity–ecosystem functioning relationships.

## Discussion

Here, we present a method to disentangle multiple components of the Net Biodiversity Effect (NBE) on functions other than primary productivity or total biomass. In essence, the method quantifies the extent to which a community functions differently than the sum of its isolated, constituent species. This method assumes that each species contribution to a function varies proportionally with its biomass. Thus, starting from the measurement of some ecosystem-level variable *V* (e.g. soil N concentration), the first step of our method is to define an associated biomass-scalable function *Φ*, proportional to community biomass, that determines, potentially in a non-linear way, the contribution of the community to the ecosystem variable (see Figure 1). Under the assumption of linearity, a sensible additive model for *Φ* can then be built based on measurements of *V* in monocultures. We then partition the ratio of realised to expected function *Φ* (which defines the NBE) into two components, which themselves divide in two more, capturing key pathways by which biodiversity impacts ecosystem functioning. The first group focuses on diversity effects on total biomass (representing *functional quantity*). It measures the extent to which community biomass, driven by the expression of species’ interaction and response traits, differs from what would have been expected based on monocultures. This term has two parts, called selection and complementarity effects on community biomass (CE and SE, respectively; corresponding rather closely to the effects defined in Loreau & Hector (2001). SE represents the tendency for the more productive species in isolation (high quantity) to dominate in the mixture. CE quantifies the extent to which ecological assembly is not a zero-sum game. The second, more novel, group of terms focuses on the diversity effect on the community’s *functional quality*. It measures the extent to which the mass-specific contribution of the community to the function of interest, reflecting the expression of species effect traits, differs from what would have been expected based on monocultures. This term also divides in two parts, called selection and complementarity effects on effect trait expression (denoted SETE and CETE, respectively). SETE captures the extent to which species that have a high mass-specific functional contribution in monoculture (high functional quality), end up dominating in the mixture. On the other hand, CETE captures the extent to which the mass-specific contribution of species depends on the biotic context, and thus aims at quantifying changes in species effect trait adjustments, caused by the presence of other species. Using data from a biodiversity-ecosystem functioning experiment of grassland plant species in controlled conditions, we illustrate that the importance of biodiversity effects on effect trait expression —a component of the NBE that is absent when the function is total biomass— can range from negligible for some ecosystem functions (e.g. soil hydraulic conductivity) to major for others (e.g. N retention capacity, forage digestibility).

### Toward a more comprehensive partitioning of biodiversity effects

Empirical BEF research has shifted towards the study of multiple ecosystem functions, whereas formal theory is only recently generalising beyond biomass (van der Plas, 2019; Guo et al., 2025; Orr et al., 2024). The original NBE partitioning (*sensu* Loreau & Hector 2001) into selection and complementarity effects has been intended to capture biodiversity effects on primary productivity and has been mostly used on total biomass (because in perennial grasslands aboveground biomass harvested annually corresponds to productivity). Our approach explicitly separates changes in ecosystem functions resulting from shifts in total community biomass from those arising from variation in mass-specific species contributions, while retaining the NBE structure. In a similar vein, Tatsumi and Loreau (2023) recently integrated functional traits by decomposing the classic NBE components into density-versus size-mediated processes. While their approach also aims to disentangle the effect of functional traits versus density, it remains fundamentally anchored to biomass. In addition, it supposes that mass-specific species contributions in the community is known, which may be possible for some traits (size, dry matter content), but not for most mass-specific community contributions (e.g. flammability, nutrient retention). Our formalism thus improves the study of ecosystem multi-functionality by disentangling the NBE on ecosystem functions into two separate pathways: biodiversity effects via shift in total biomass and biodiversity effect via the expression of effect traits.

The partitioning approach we present here applies specifically to biomass-scalable functions, i.e. functions to which the community contribution is proportional (linearly related) to its total biomass. Many ecosystem variables directly qualify as biomass-scalable functions (e.g. primary production, nutrient uptake, forage provisioning, or pollination), so our partitioning approach can be directly applied. However, some ecosystem variables need to be transformed in order to obtain biomass-scalable functions (see Box 1). Without an appropriate conversion, our partitioning may detect large artifactual biodiversity effects. Because the NBE captures the departure of an observation from an expectation, any flaw in the expectation will be reflected in the resulting NBE. For example, if the variable ‘N concentration of leachates’ had not been properly standardised and transformed to a biomass-scalable function, we would have obtained high biodiversity effects (see Appendix A). The transformation to biomass-scalable functions has to be driven by a mechanistic understanding of the processes underlying the measured ecosystem variables. Finding a generic way to transform ecosystem variables to biomass-scalable functions, in a way that integrates non-linearity (in the vein of Baert et al., 2017), thus requires that a choice is made regarding the shape of the relationship between the ecosystem variable and the biomass-scalable function. This choice can be based on measurements of the ecosystem variable against a (controlled) biomass gradient, e.g. during a growth phase.

### Revealing the signature of trait expression in biodiversity effects

Our partitioning framework, though descriptive, aims to shed light on the mechanisms underlying community effects on a wide range of ecosystem variables. Here, we discuss the effect of plants on several biomass-scalable functions within the short time scale (15-month) experiment of Bittlingmaier et al. (2026) to illustrate the interpretative value of this partitioning. Following a theoretical development (Box 1), we assumed that the bulk of plants’ contribution to ecosystem N retention is due to their effective N uptake capacity. We therefore neglected the indirect effect of plants on soil microbial communities and abiotic properties influencing N retention which would unfold at longer timescales. We observed a negative selection effect on effect trait expression, most plausibly explained by the dominance of N-fixing legumes in polycultures (Temperton et al., 2006). Because legumes can rely on symbiotically fixed N after early reliance on soil N uptake, they are able to produce more biomass from a given concentration of soil N (Vitousek et al., 2002), resulting in lower per-biomass uptake rates. the negative complementarity effect on trait expression observed for N retention capacity suggests that, in species-rich communities, community-level plant plastic adjustments tend to further decrease their N uptake rates. This negative CETE may reflect the phenotypic plasticity of some species that are able to resort to other types of resources (e.g. ammonium vs. nitrate, Ardichvili et al., 2024), or from biomass allocation strategies (roots vs. stems) that differ between the intra and inter-specific contexts (Rehling et al., 2021). The positive complementarity effect on trait expression for forage digestibility likely reflects phenotypic adjustments driven by light competition. In dominant species, stronger intra-specific competition for light may lead to greater investment in plant structural tissues which typically have high dry matter content and are less digestible (Rehling et al., 2021). This shade-avoidance response could cause these species to grow higher and reach more light (Violle et al., 2009). The improvement of soil hydraulic conductivity with higher species richness can be attributed solely to increases in total biomass. Having more biomass in a polyculture may result in a more permeable soil structure due to higher root density leaving more channels open for water penetration after root turnover (Hao et al., 2020). The absence of biodiversity effects on effect trait expression indicates root turnover rates (or any other functional traits considered important for soil hydraulic conductivity) may not differ strongly between species, or that there is no specific synergistic effect on trait expression due to higher plant diversity.

Understanding the functional role of biodiversity requires considering its effects on both species traits and total biomass. Approaches that focus on species traits alone, even weighted by relative biomass, provide only a partial view of how biodiversity affects ecosystem functioning. Likewise, the classical additive partitioning of selection and complementarity is insufficient because it conflates trait-mediated and biomass-driven effects on broader ecosystem functioning. By contrast, the four components of NBE presented here represent these signatures of species interactions, expressed through both biomass and effect trait expressions, thus their effect on quantity *and* quality. Since the underlying mechanisms remain hypothetical, this method could be further strengthened by coupling it with trait measurements. In particular, refining or confirming our interpretations would require linking changes in mass-specific contributions to functional traits measured at the species or community level (e.g. root length density and root turnover rate for the soil hydraulic conductivity function). In this vein, Michel et al. (2012) computed biodiversity effects on water retention in bryophyte communities and showed that some traits (cushion height and shoot area-to-volume ratio) were correlated with water retention, whereas other traits (shoot density) were associated with the observed biodiversity effects. Traits that only affect the ecosystem function (cushion heights and shoot area-to-volume ratio in the study of Michel et al., 2012) are arguably closely linked to the mass-specific contribution of species. Traits that are linked to biodiversity effects (shoot density) are traits which are indirectly linked to the function, but operate by mediating interactions between species. While explicitly linking functional traits to mass-specific species contributions (Fig. 1) is lacking from both our approach and the one of Michel et al. (2012), it represents an important next step for understanding how biodiversity, functional traits, and total biomass shape ecosystem functioning.

## Conclusion

The biodiversity effects we measure in experiments indicate when the functional impact of a species is dependent on the biotic context in which it grows. As such, biodiversity effects are signatures of biotic interactions. While it may seem trivial to state that biodiversity effects on total biomass reflect species interactions, it is worth noting that the effect on a species can go beyond a change in its biomass. The presence of other species can cause phenotypic adjustments, which will in turn leave a signature in the biodiversity effects on effect trait expression. Recognising this dual role of biodiversity is crucial for effectively conserving the eroding services biodiversity provides (Violle et al., 2007).

## Acknowledgment

This work was supported by the BIOSTASES Advanced Grant, funded by the European Research Council (ERC) under the European Union’s Horizon 2020 research and innovation programme (666971), the Laboratoire d’Excellence TULIP (ANR-10-LABX-0041), the École Universitaire de Recherche TULIPGS (ANR-18-EURE-0019), the ANR project REFRAME (Grant ANR-24-CE02-7222), and the Occitanie Region.

## Appendix A Example of artifactual biodiversity effects on trait expression

We continue to analyse the model of N cycling developed in Box 1. We consider two species, A and B, that reach biomass levels K_A_ and K_B_ in isolation and B_A_ and B_B_ in duo-culture, when seeded at equal densities. The goal of this appendix is to illustrate the importance of properly identifying the biomass-scalable function associated to an ecosystem variable. To do so, we here purposely omit to distinguish the function from the ecosystem variable measured (inorganic soil N concentration), and derive artifactual biodiversity effects on trait expression caused by this omission.

1. *Measuring the ecosystem function in monoculture:* In monoculture, soil inorganic N concentration at equilibrium reads:

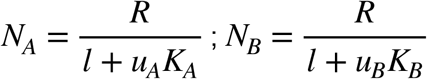
2. *Measuring observed soil N in the mixture:* In a mixture of the two species, assuming fixed uptake rates (so no change in effect traits), soil inorganic N concentration at equilibrium reads: :

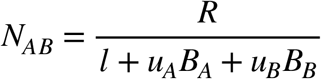

We see that this quantity is not proportional to total biomass and therefore is not a biomass-scalable function. If we nonetheless treat it as such we would compute a community mass specific contribution as:

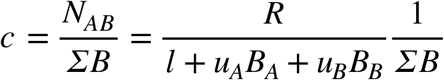
3. *Deducing expected mass-specific contributions from monocultures:* If soil N concentration was (wrongly) treated as a biomass-scalable function, we would conclude that the expected mass-specific contribution of each species is:

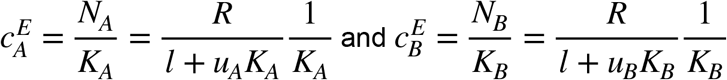

Given species realised biomass we would then build the expected soil N concentration as:

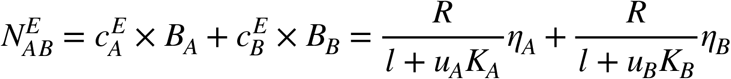

Where *η*_*A,B*_ stands for the species relative yield. The expected mass-specific community contribution becomes:

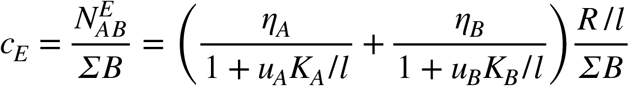
4. *Computing biodiversity effects* For simplicity we assume that in the mixture, the two species reach the same relative yield *η* This means that we assume no selection effects on trait expression (relative densities are as expected from monocultures). The two components of the CETE, c and c_E_ then read:

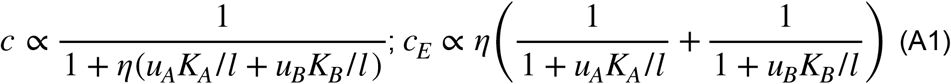

where the common proportionality constant is 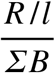 . From Eq. A1 (and Fig. A1), we see that c and c_E_ go in opposite directions as *η* increases. c=c_E_ for only one value, and this value depends on the various nutrient cycling parameters. This means that, although the model does not contain any variations in species effect traits, we may still obtain a CETE larger than 1, smaller than 1, or equal to 1 depending on the context, but unrelated to an actual trait variation. This demonstrates that a CETE that strongly differs from 1 may signal that what is used to obtain mass-specific contributions may not have been appropriately transformed to a biomass-scalable function.

**Figure A1.**
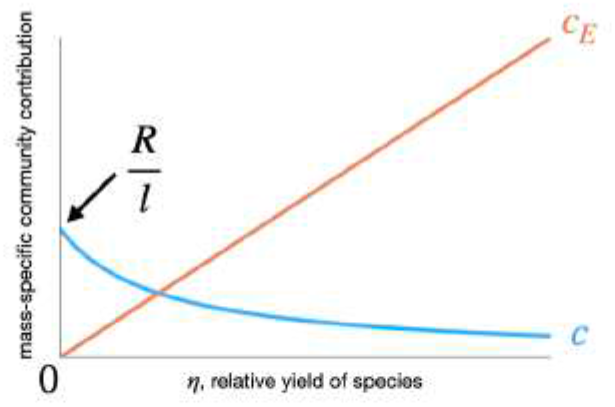
expected versus realised mass-specific community contribution as a function of the relative yield of the two species.

